# scRNA-Sequencing uncovers a TCF-4-dependent transcription factor network regulating commissure development

**DOI:** 10.1101/2020.06.16.156083

**Authors:** Marie-Theres Wittmann, Philipp Kirchner, Arif B. Ekici, Elisabeth Sock, D. Chichung Lie, André Reis

## Abstract

Intercortical connectivity is important for higher cognitive brain functions by providing the basis for integrating information from both hemispheres. We show that ablation of the neurodevelopmental disorder associated bHLH factor *Tcf4* results in complete loss of forebrain commissural systems in mice. Applying a new bioinformatic strategy integrating transcription factor expression levels and regulon activities from single cell RNA-sequencing data predicted a TCF-4 interacting transcription factor network in intercortical projection neurons regulating commissure formation. This network comprises a number of regulators previously linked to the pathogenesis of intellectual disability, autism-spectrum disorders and schizophrenia, e.g. *Foxg1, Sox11* and *Brg1*. Furthermore, we demonstrate that TCF-4 and SOX11 biochemically interact and cooperatively control commissure formation *in vivo*, and regulate the transcription of genes implied in this process. Our study provides a regulatory transcriptional network for the development of interhemispheric connectivity with potential pathophysiological relevance in neurodevelopmental disorders.

## Introduction

Cognitive abilities are highly dependent on the establishment of proper neuronal connectivity between different brain regions and its cellular components (Constantinidis and Klingberg 2016; Hedden and Gabrieli 2004). The corpus callosum, the anterior commissure, and the hippocampal commissure, carry axons across the midline and ensure information flow and coordination between the cerebral hemispheres. Of these, the corpus callosum (CC) is the largest commissural tract of the human brain (Tomasch 1954). Callosal connections serve to integrate and coordinate of sensory-motor functions from the right and left side of the body and are integral to high-level cognitive functions including language, abstract reasoning and high-level associative function (Paul et al. 2007).

Mutations in a number of factors controlling developmental processes such as neuronal precursor proliferation, fate specification, migration and axon guidance, are associated with structural anomalies of commissures (Edwards et al. 2014; Lindwall et al. 2007; Paul et al. 2007; Richards et al. 2004), illustrating that commissure development is highly dependent on the expression of complex genetic programs. Orchestration of the precise temporo-spatial execution of developmental programs is most likely achieved by cell-type specific combinatorial activity of transcription factors. However, information on the composition of transcription factor networks in commissural development remains scarce and is largely confined to the regulation of upper layer neuron specification, which is highly reliant on a SATB2-dependent genetic network (Alcamo et al. 2008; Britanova et al. 2008).

The class I basic helix-loop-helix (bHLH) transcription factor (TF) TCF-4 (transcription factor 4) has recently emerged as a critical transcriptional regulator in forebrain development. Variants in *TCF4* have been associated with schizophrenia, autism and intellectual disability and *TCF4* haploinsufficiency causes the neurodevelopmental disorder Pitt-Hopkins syndrome (PTHS) (OMIM 610954) (Amiel et al. 2007; De Rubeis et al. 2014; Navarrete et al. 2013; Schizophrenia Psychiatric Genome-Wide Association Study 2011; Schizophrenia Working Group of the Psychiatric Genomics 2014; Stefansson et al. 2009; Steinberg et al. 2011; Zweier et al. 2007).

Alterations of TCF-4 dosage in mice lead to disruptions in neocortical neuronal migration, specification of neuronal subtypes, dendrite and synapse formation (Li et al. 2019; Page et al. 2017). Most notably, *TCF4* haploinsufficiency in human and mice is associated with callosal dysgenesis indicating that *Tcf4* is part of a conserved genetic network controlling the formation of callosal connections (Jung et al. 2018).

TCF-4 belongs to the class I basic basic Helix-Loop-Helix (bHLH) transcription factor family and its transcriptional output is highly dependent on its interaction partners. Traditionally, it is assumed that TCF-4 executes its function through dimerization with proneural class II bHLH TFs (Bertrand et al. 2002; Massari and Murre 2000). A recent *in vitro* study, however, proposed that TCF-4 may be able to interact with a variety of transcriptional regulators outside of the traditional interaction partners of the bHLH class (Moen et al. 2017). TCF-4-interacting transcription factors in the regulation of interhemispheric connectivity have not been identified. Such identification is hampered by the technical challenges to perform unbiased in vivo interactome analyses in a cell type specific manner.

Here, we generated homozygote *Tcf4* knockout (*Tcf4KO*) mice to further validate the role of TCF-4 in the establishment of interhemispheric connectivity. We report that loss of *Tcf4* results in the complete agenesis of forebrain commissures. Using single-cell RNA sequencing (scRNA-seq) followed by the integration of transcription factor expression levels and regulon activities we uncover a TCF-4 interacting transcription factor network for commissure development in *Satb2* expressing neurons. Surprisingly, this transcription factor network involves numerous transcription factors outside the bHLH class. Similar to TCF-4, these interactors are often associated to neurodevelopmental disorders such as intellectual disability, autism and schizophrenia. Further analysis of the interaction between TCF-4 and the regulator SOX11 uncovered a synergistic effect on anterior commissure (AC) and CC formation. Collectively, our findings provide insight into a novel gene regulatory network controlling commissure formation and potentially involved as a common pathogenic pathway in neurodevelopmental and neuropsychiatric diseases.

## Results

### *Tcf4* knockout abolishes commissure development

*Tcf4* haploinsufficient mice generated by an insertion of a lacZ/neomycin cassette before exon 4 (Figure 1A) show dysgenesis of the corpus callosum (Jung et al. 2018). To further examine the critical role of *Tcf4* in development of interhemispheric connectivity, we generated *Tcf4* homozygote knockout mice. Effectiveness of the *Tcf4* knockout was assessed by western blot as there are *Tcf4* isoforms with transcriptional start sites after exon 4 (Sepp et al. 2011). Our analysis confirmed the loss of the longest TCF-4 isoform (TCF-4B) in the knockout; expression of a shorter isoform (TCF-4A) persisted (Figure 1B). Despite the residual TCF-4 expression, *Tcf4KO* mice died shortly after birth, indicating the importance of the long TCF-4 isoform during development.

**Figure 1.**
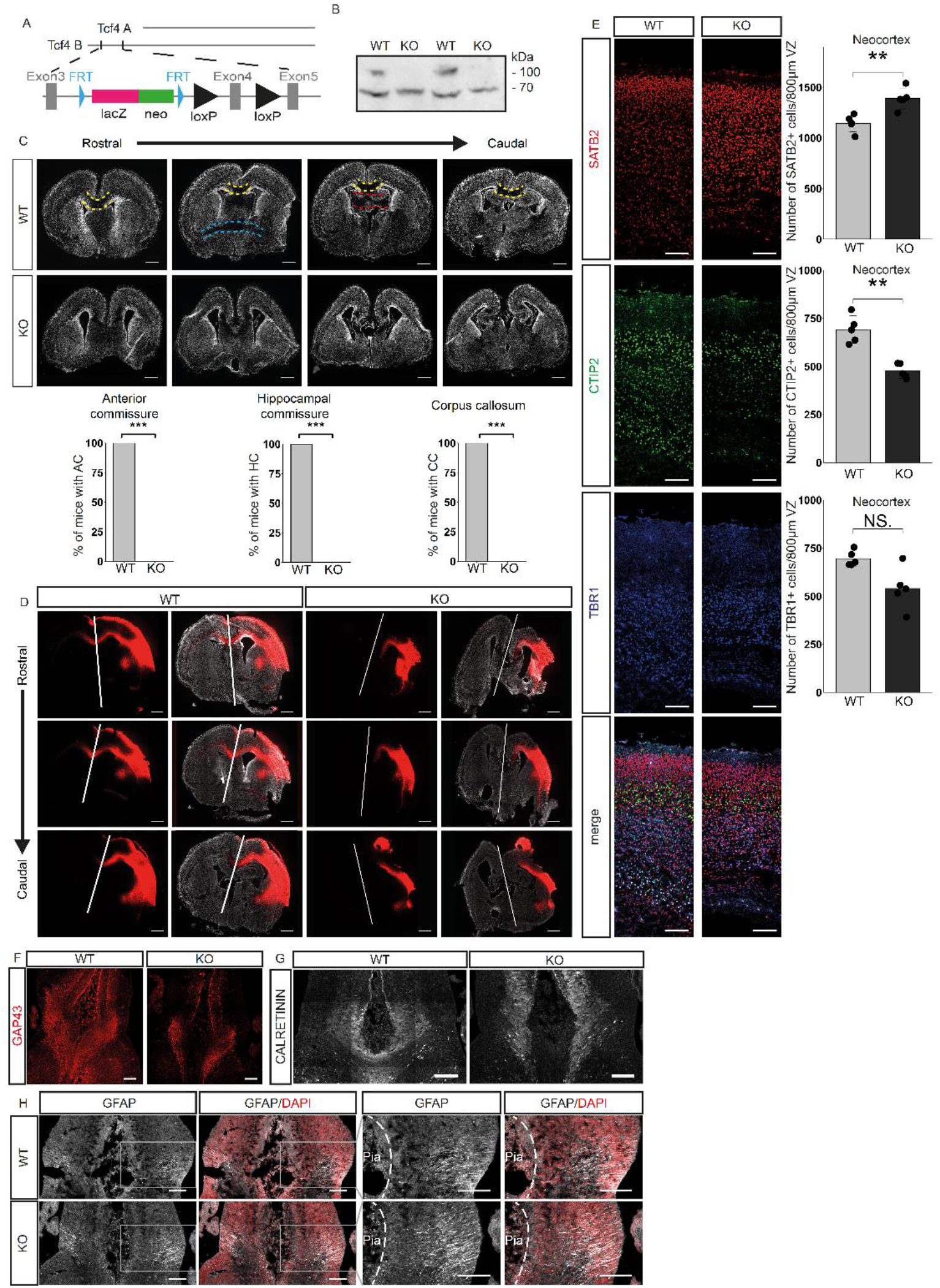
Loss of *Tcf4* disrupts cortex development, especially commissure formation. **A** Schematic representation of the two main *Tcf4* isoform and the ‘knockout-first’ conditional allele. **B** Western Blot analysis of neocortical extracts from E18.5 WT or *Tcf4KO* mice using anti-TCF-4 antibody. The blot presented is cropped. The longest isoform of TCF-4 is missing in the KO samples (n = 3). **C** Representative overview images (DAPI) of brain sections at P0 showing the loss of the three commissure systems in *Tcf4KO* mice. Yellow dotted lines indicate the CC crossing the midline. Blue dotted lines indicate the AC and red dotted lines the HC. Quantification of animals showing a commissural system is presented below (n = 8). Scale bar, 500 μm. Statistical significance was determined by two-tailed Mann-Whitney-U test (*, P ≤0.05; **, P ≤0.01, ***, P ≤0.001) **D** Representative images of lipophilic tracer (red) treated brains at P0 without or with DAPI staining (white). White lines indicate the midline. Note that only in WT animals, lipophilic tracer signal can be detected in the contralateral hemispheres to the treatment (n = 3). Scale bar, 500 μm. **E** Representative images of the neuronal markers SATB2 (upper layers), CTIP2 (layer V) and TBR1 (layer 6) and the quantification of the total cell number expressing these markers per 800μm ventricular zone (VZ). A significant increase in SATB2+ neurons is observable as is a significant decrease in CTIP2+ cells. Scale bar 100μm. Data is presented as mean ± SD; (n = 5). Statistical significance was determined by two-tailed Mann-Whitney-U test (*, P ≤0.05; **, P ≤0.01, ***, P ≤0.001). **F** Representative images for GAP43 at the midline of E16.5 mouse brains. Scale bar, 100 μm, (n = 3). **G** Representative images of CALRETININ expression at the midline at E16.5. Scale bar, 100 μm, (n = 3). **H** Representative images of GFAP stainings at E16.5. Pictures on the right side are magnifications of the marked area. Dotted lines represent the pial surface. Scale bar, 100 μm, (n = 3).

The most striking feature of P0 KO brains was the absence of the forebrain commissure system, i.e. CC, AC, and hippocampal commissure (Figure 1C, D). Staining for the upper layer and interhemispheric projection neuron marker SATB2 (Alcamo et al. 2008; Britanova et al. 2008) revealed a significant increase in SATB2 positive neurons (SATB2+ cells/800μm VZ: WT 1146 ± 75.89; KO 1389 ± 92.45; p = 0.0079), which appeared to be generated at the expense of CTIP2 (CTIP2+ /800μm VZ: WT 691 ± 64.2; KO 476 ± 33.9; p = 0.0079) but not of TBR1 positive deep layer neurons (TBR1+ cells/800μm VZ: WT 695 ± 35.5; KO 540 ± 97.7; p = 0,0556; Figure 1E). These data indicate that the commissure forming SATB2+ neurons were in principle generated in *Tcf4KO* mice. Analysis with the anterograde lipohilic tracer DiI indicated that neurons in *Tcf4KO* mice extended their axons towards the midline yet were unable to cross to the contralateral side (Figure 1D). Staining for GAP43, a marker for the axonal growth cone verified this observation (Figure 1F).

Callosal agenesis can be the consequence of dysfunctional midline generation and fusion (Richards et al. 2004). We analysed mice at E16.5, to investigate the characteristic detachment of extensions to the pial surface by GFAP-expressing glial wedge glia and the presence of Calretinin-expressing guidepost neurons (Paul et al. 2007). At E16.5 both cell types were detected at the midline in *Tcf4KO* mice and no general defect in the organization of the structure was observed (Figure 1G and H), suggesting that the midline had been properly formed.

### *Tcf4* knockout dysregulates genes involved in neuronal differentiation and axon guidance

The presence of SATB2 positive neurons and the formation of a midline in *Tcf4KO* mice raised the possibility that TCF-4 may regulate in particular the formation of the axonal tracts by SATB2 expressing neurons. To identify downstream targets and pathways of TCF-4 in the control of forebrain commissure formation, scRNA-seq was conducted from E18.5 neocortices. After rigorous filtering for viable cells, 8,887 cells were analysed for the WT and 5,309 for the KO. Cell clustering was performed using Seurat and cell types were assigned using known markers (Figure 2A) (Butler et al. 2018; Stuart et al. 2019). All expected major cell types were detected and WT and KO cells clustered together regardless of their genotype (Figure 2B).

**Figure 2.**
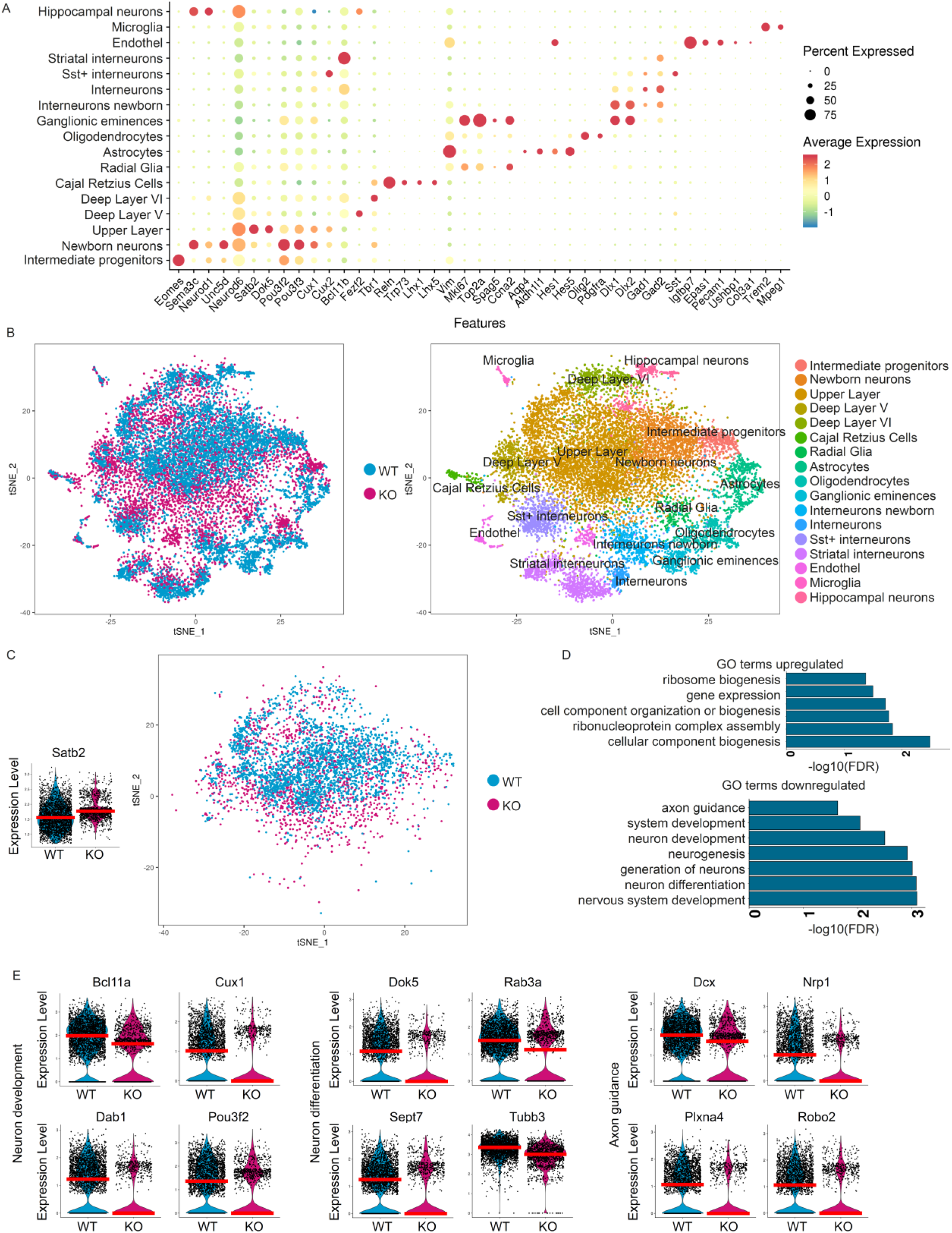
Single-cell RNA Sequencing of E18.5 neocortices from WT and *Tcf4KO* mice. **A** Dot-Plot of cell clusters (y-axis) and marker used to assign the cell type (x-axis). **B** tSNE-Plot coloured by genotype (left) and cluster identity (right). **C** tSNE-Plot of *Satb2* expressing glutamatergic cells used for further analysis. Violin Plot of *Satb2* expression in the *Satb2* cluster. The red line depicts the median. **D** GO terms associated with up- and downregulated genes in SATB2 expressing glutamatergic cells. GO terms for neurogenesis, neuronal differentiation and axonogenesis were downregulated in the *Tcf4*KO cells. **E** Violin Plot of differentially expressed genes in the *Satb2* cluster that are associated to neuron development/differentiation and axon guidance. The red line depicts the median.

To define TCF-4-dependent networks in commissure forming neurons, the MAST algorithm was used to determine differentially expressed genes (DEGs) between *Satb2* expressing glutamatergic WT (2,890 cells) and KO (1,328 cells) neurons (Figure 2C and Table S1) (Finak et al. 2015). Upregulated genes (97 genes) were associated with GO terms for ribosomes and gene expression (Figure 2D). GO terms for downregulated genes (131 genes) included gene enrichments for neuron development (e.g. *Bcl11a, Cux1, Dab1 and Pou3f2*) and differentiation (e.g. *Dok5, Rab3a, Sept7 and Tubb3*) as well as axon guidance (e.g. *Dcx, Nrp1, Plxna4, and Robo2*) (Figure 2D and E).

Analysis of unique molecular identifiers (UMIs), number of genes and the percentage of mitochondrial genes revealed that KO cells had in general a lower number of expressed genes and UMIs. We therefore limited the data to cells with less than 4000 UMIs, yielding 1057 WT and 1246 KO cells and re-analyzed DEG and GO term enrichment to account for potential bias introduced by differences in gene and cell number in the *Satb2* cluster. Most DEGs and GO terms remained the same indicating the biological relevance of our findings (Table S2).

Collectively, these findings molecularly confirm the anatomical observation that *Tcf4* knockout affects the expression of a gene network in *Satb2* neurons, that is involved in commissure and neuron projection development.

### TCF-4 modulates the regulon activity of neurodevelopmental transcription factors

We next aimed to investigate the influence of TCF-4 on gene regulatory networks (GRNs). GRNs were calculated using the R package of SCENIC (Aibar et al. 2017). Identified regulons were binarized to generate an ON/OFF state (Figure 3A). Re-clustering displayed a partitioning of cells between the two genotypes with only marginal overlap (Figure 3A). Regulons were sorted into three categories: In the first, the regulon was primarily active in the WT. In the second, there was no differential activity between the genotypes, and in the third, the regulon was predominantly active in the KO. Differentially active regulons mostly fell into the first category [e.g. *Ctcf*, *Foxg1, Smarca4* (also known as *Brg1*), *Cux1*, *Pou3f3* (also known as *Brn1*), and *Sox11]* (Figure 3B and C and Table S2). As with the DEG analysis, we repeated the analysis with the limited *Satb2* dataset and replicated these findings (Table S3). Further analysis was therefore done with the original dataset.

**Figure 3.**
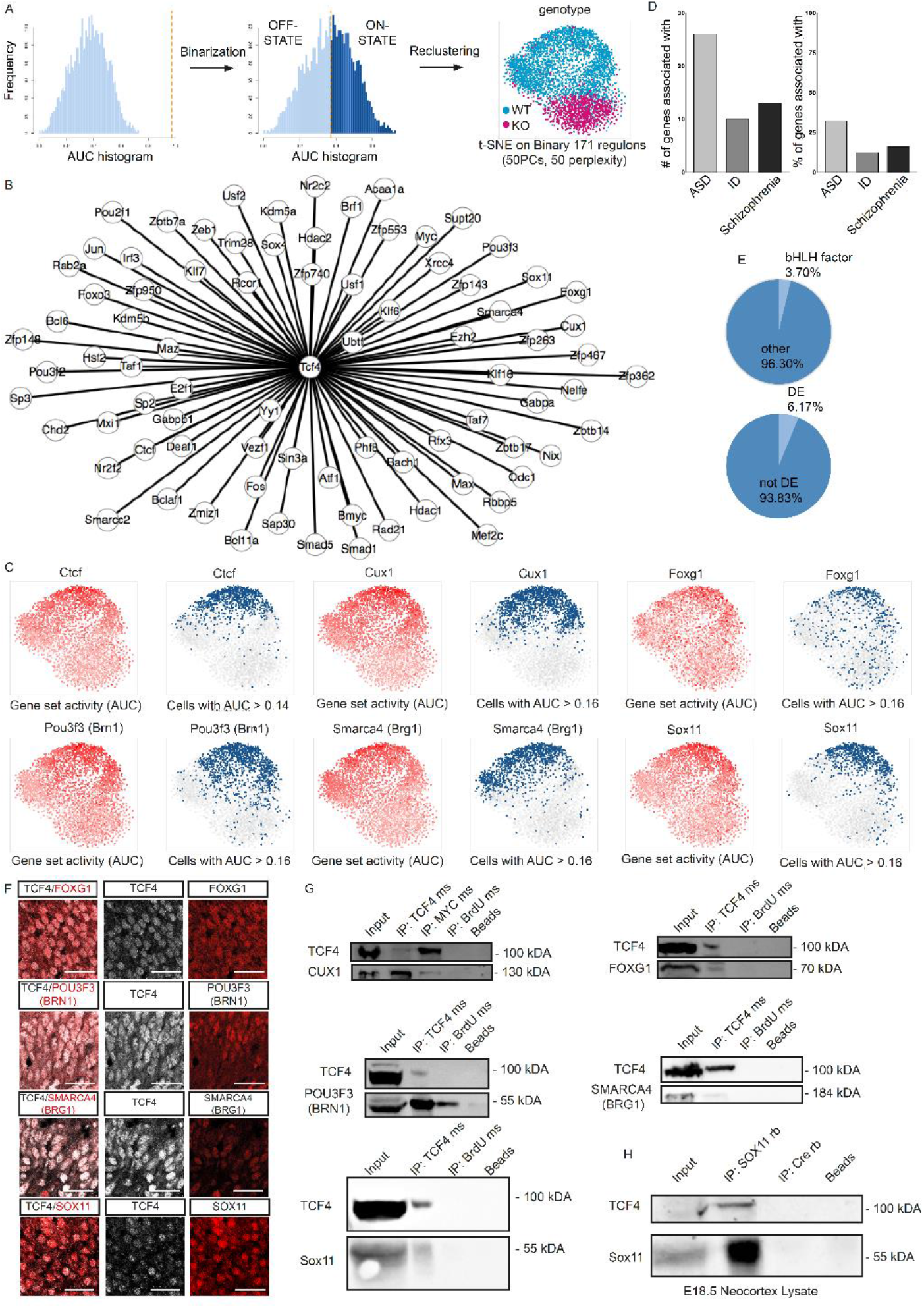
Gene regulatory network analysis of SATB2 expressing cells. **A** Scheme of the workflow used to recluster cells after GRN analysis and resulting tSNE-Plot of the *Satb2* cluster. Regulons are binarized and reclustered accordingly. WT and KO cells segregated based on GRN activity with only minor overlap. **B** Differentially active regulons of the *Satb2* cluster that may be possible interactors of TCF-4. **C** tSNE-Plots showing the regulon activity of *Ctcf, Cux1, Foxg1, Pou3f3* (also known as *Brn1*), *Smarca4* (also known as *Brg1*) and *Sox11* in a continuous scale (left, red) or binarized (right, blue). The regulons are preferentially active in the WT cells with only a small number of KO cells in the ON-State. **D** Number and percentage of regulators associated with autism spectrum disorders (ASD), intellectual disability (ID) and schizophrenia in the *Satb2* cluster. **E** Pie charts depicting the percentage of bHLH factors and differentially expressed regulators in the differentially active regulons. **F** Representative images of TCF-4 (white) and FOXG1, POU3F3 (BRN1), SMARCA4 (BRG1) and SOX11 (all in red) in E18.5 WT cortices. Note the expression in the same nuclei. Scale bar, 50 μm **D** Co-immunoprecipitation assay using anti-TCF-4 antibody conducted with HEK cell extract after overexpression of TCF-4 and CUX1, FOXG1, POU3F3 (BRN1), SMARCA4 (BRG1) and SOX11 in HEK cells. Upper panels: detection with anti-TCF-4 antibody. Lower panels: detection with anti-MYC, anti-FOXG1, anti-BRN1, anti-BRG1 or anti-SOX11 antibody. The blots presented are cropped. All proteins were co-immunoprecipitated with TCF-4, but not with an isotype control for IgG or Agarose A Beads alone except for BRN-1 which was precipitated to a small amount by the isotope control IgG. We could also show that TCF-4 was co-immunoprecipitated with anti-Myc antibody (precipitation of Myc-tagged Cux1). The interactions were confirmed in three independent biological replicates, (n = 3). **E** Co-immunoprecipitation assay conducted with E18.5 cortex lysates using anti-SOX11 antibody. Upper panel: detection with anti-TCF-4 antibody. Lower panel: detection with anti-SOX11 antibody. The blots presented are cropped. TCF-4 was co-immunoprecipitated with SOX11, but not with an isotype control for IgG and Agarose A Beads alone. The interaction was confirmed in three independent biological replicates (n = 3).

Examination of the differentially active regulons revealed that most regulon heads functioned as TFs. Interestingly, the respective genes were to a high extent associated with autism spectrum disorders (ASD), intellectual disability (ID) and schizophrenia [ASD: 26 genes; 32.10%; e.g. *FOXG1*; ID: 10 genes; 12.35%; e.g. *CTCF*; Schizophrenia: 13 genes; 16.05%; e.g. *SOX11* (Figure 3D)](Gregor et al. 2013; Kortum et al. 2011; Sun et al. 2020). To our surprise, these TFs generally did not belong to the bHLH TF family, which constitutes the canonical interaction partners of TCF-4 (bHLH factors: 3.70%; Other: 96.30%) (Figure 3E). In addition, the vast majority of regulators did not appear to be downstream targets of TCF-4 as they were mostly not differentially expressed in the KO (DE: 6.17%; Not DE: 93.83%) (Figure 3E). This observation raised the question of how the loss of TCF-4 impacted on regulon activity. A recent *in vitro* study suggested that TCF-4 may interact with TFs outside the bHLH family (Moen et al. 2017), leading to the hypothesis that TCF-4 interacts with the regulators and thereby modulates their activity. We thus focused on potential interactions regulating genes involved in neurogenesis or neuron differentiation. To ensure a focus on the robustly expressed regulons, we required the selected regulators to be expressed in at least one quarter of the *Satb2* expressing cells (Table S4). Five regulon heads [*Foxg1, Smarca4* (also known as *Brg1*), *Cux1*, *Pou3f3* (also known as *Brn1*), and *Sox11*] were selected for validation (Figure 3C). CUX1 and POU3F3 are specific markers for layer II/III neurons, whereas FOXG1, SMARCA4 and SOX11 are broadly expressed during neuronal differentiation (Bergsland et al. 2006; Campbell et al. 2008; Deng et al. 2015; Miyoshi and Fishell 2012; Molyneaux et al. 2007; Seo 2004). Staining of neocortical tissue at E18.5, showed that TCF-4 was co-expressed with SOX11, FOXG1, SMARCA4 and POU3F3 (Figure 3F). *In vitro* co-immunoprecipitation assays confirmed that the long TCF-4 isoform has the potential to biochemically interact with all these TFs (Figure 3G). Moreover, co-immunoprecipitation assays from E18.5 neocortex lysates validated the interaction of TCF-4 with SOX11 *in vivo* (Figure 3H).

Thus, TCF-4 has the ability to biochemically interact with a wide variety of TFs and chromatin remodelers involved in neurogenesis and neuronal differentiation and may thereby modulate their activity during commissure development.

### TCF-4 and SOX11 act synergistically during commissural development

We focused our successive investigation on the interaction of TCF-4 and SOX11. SOX11 contributes to an evolutionary conserved program controlling axonal tract formation of corticospinal neurons (Shim et al. 2012). SOX11’s importance for human neurodevelopment is demonstrated by its causal link to a Coffin-Siris-like syndrome (OMIM 615886), a neurodevelopmental disease characterised by microcephaly and intellectual disability (Hempel et al. 2016; Turan et al. 2019). Assessing the mRNA levels in the *Satb2* cluster, *Sox11* was found to be expressed in almost every cell, rendering it the most commonly expressed regulator examined.

To evaluate the functional interaction of TCF-4 and SOX11 *in vivo, Tcf4* and *Sox11* haploinsufficient mice were crossed to generate WT, *Tcf4, Sox11* and double *Tcf4* and *Sox11* haploinsufficient littermates. Commissural systems in P56 brains were visualized by Luxol fast blue staining. WT and *Sox11* haploinsufficient mice showed no commissural phenotype (Figure 4A). In line with a previous report, *Tcf4* haploinsufficient animals had a mildly shortened CC (data not shown) (Jung et al. 2018). This phenotype was greatly aggravated by the additional haploinsufficiency of *Sox11* as double haploinsufficient mice showed the most severe truncation of the CC with only the most rostral part of the CC remaining (agenesis of the splenium and caudal part of the body) (Figure 4A). Moreover, only a rudimentary AC (1 out of 5 animals) or a complete agenesis of the AC (4 out 5 animals) was observed in the double haploinsufficient mice (Figure 4A).

**Figure 4.**
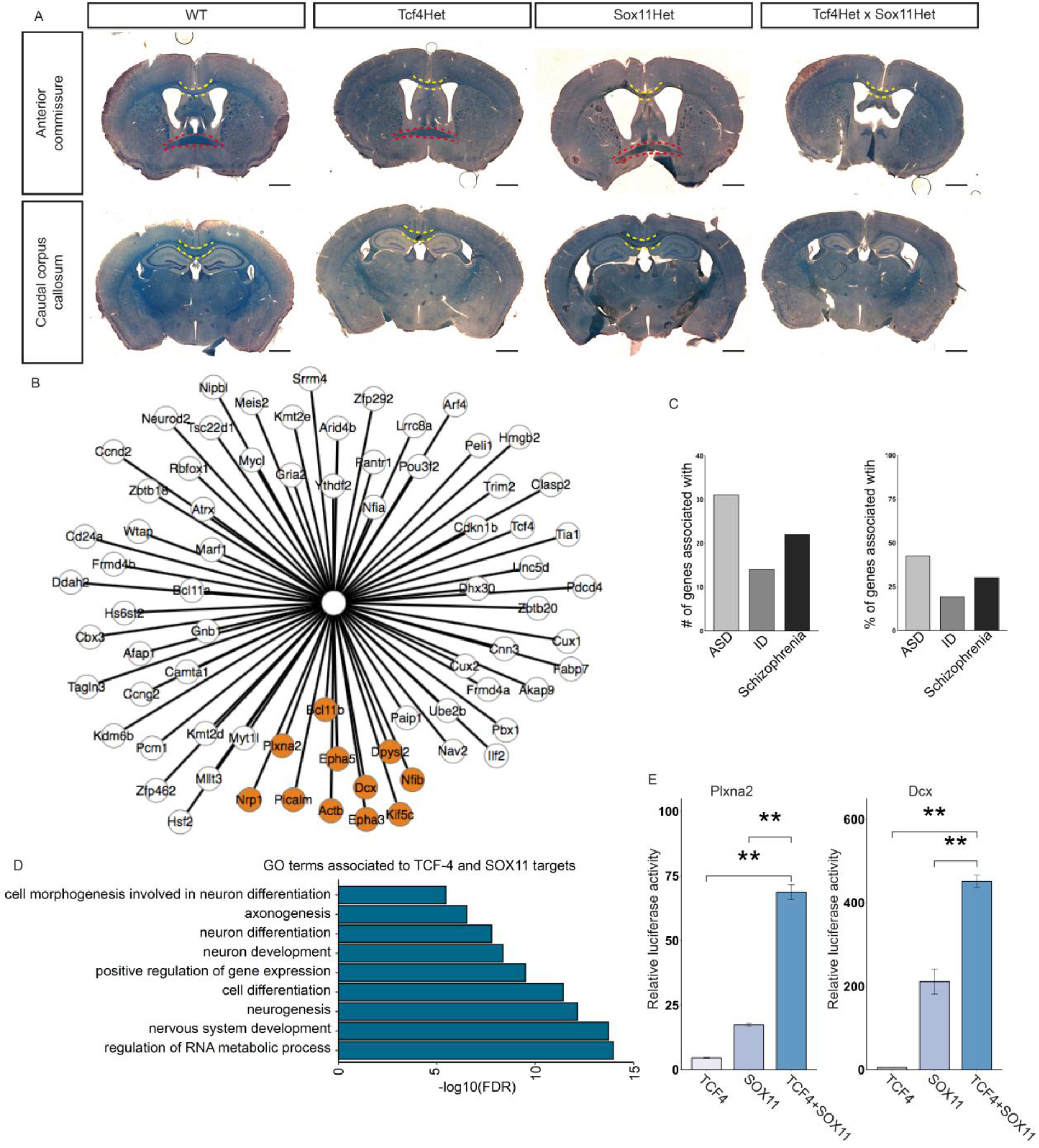
TCF-4 and SOX11 act synergistically in corpus callosum formation. **A** Representative overview images of Luxol fast blue stainings at the position of the AC and the caudal body of the CC. Yellow dotted lines indicate the CC crossing the midline. Red dotted lines indicate the AC. In *Tcf4* and *Sox11* double haploinsufficient mice agenesis of the AC and agenesis of the splenium and caudal part of the body of the CC can be observed. Scale bar, 1000 μm, (n=5). **B** Common targets of TCF-4 and SOX11 in the *Satb2* cluster. Orange highlighted genes are associated with the GO term axonogenesis.**C** Number and percentage of common targets of TCF-4 and SOX11 in the *Satb2* cluster associated to autism spectrum disorders (ASD), intellectual disability (ID) and schizophrenia in the *Satb2* cluster.**D** Selection of GO terms associated with the common targets of TCF-4 and SOX11 in the *Satb2* cluster. GO terms for neurogenesis, neuronal differentiation and axonogenesis were enriched. **E** Relative luciferase reporter gene activity under the control of the regulatory regions from the *Plxna2* (C) and *Dcx* (D) in transiently transfected HEK cells co-expressing TCF-4, SOX11 and a combination of the two (n = 3, presented as fold induction ± SD, transfection with empty CAG-GFP vector was set to 1 for each regulatory region). Statistical significance was determined by a two-tailed student’s t-test. (*, P ≤0.05; **, P ≤0.01, ***, P ≤0.001)

We next asked which genes may be common target genes of both TFs and thus may be involved in commissure development. Hence, we compared the predicted regulon targets of SOX11 from the GRN analysis and the list of DEGs in the *Tcf4KO*. The intersection of the datasets yielded a list of 73 genes (Figure 4B and Table S3), which similarly to the differentially active regulons were often associated with autism spectrum disorders, intellectual disability and schizophrenia [ASD: 31 genes; 42.47%; e.g. *GRIA2*; ID: 14 genes; 19.18%; e.g. *DCX*; Schizophrenia: 22 genes; 30.14%; e.g. *PLXNA2* (Figure 4C)] (Mah et al. 2006; Pilz et al. 1998; Salpietro et al. 2019). Furthermore, GO term analysis revealed an enrichment for genes involved in axonogenesis (Figure 4D and Table S3). From these associated genes *Plxna2*, a gene involved in semaphorin plexin signalling for axon guidance (Mah et al. 2006; Mitsogiannis et al. 2017; Rohm et al. 2000) and *Dcx*, a gene essential for proper neuronal morphology, migration and axon guidance were selected for further investigation (Deuel et al. 2006; Fu et al. 2013; Karl et al. 2005; Koizumi et al. 2006). Evolutionary conserved regions upstream of or at the promotor, which contained conserved binding sites for TCF-4 and SOX11, were cloned into luciferase reporter plasmids and then transfected into HEK293T cells together with expression plasmids for *Tcf4* and *Sox11*. SOX11 alone induced robust *Dcx* and *Plxna2* reporter activity. TCF-4 alone only marginally induced *Plxna2* and *Dcx* activity but strongly potentiated SOX11 induced reporter activities (*Dcx*: SOX11 vs. TCF4+SOX11: p-value = 0.0059; TCF4 vs. TCF4+SOX11: p-value = 0.0011; *Plxna2*: SOX11 vs. TCF4+SOX11: p-value = 0.0018; TCF4 vs. TCF4+SOX11: p-value = 0.0021) (Figure 4B and C). Collectively, these results indicate the cooperative interaction of TCF-4 and SOX11 in AC and CC formation by activating gene expression and their importance in axonogenesis and axon guidance.

## Discussion

Interhemispheric connections are central for higher brain function by integrating information from both hemispheres (Constantinidis and Klingberg 2016; Hedden and Gabrieli 2004). Here, we show that *Tcf4* knockout severely disrupts cortex development, especially commissure formation. We provide scRNA-Seq and biochemical evidence that positions the bHLH transcription factor TCF-4 at the centre of a large regulatory network for forebrain commissure formation. Of particular interest is the finding that in this network TCF-4 interacts with multiple intellectual disability, autism and schizophrenia associated transcriptional regulators raising the possibility that the TCF-4 dependent regulatory network in commissure formation may be relevant for the pathogenesis of neurodevelopmental and -psychiatric disorders.

Previous analysis revealed the existence of multiple TCF-4 isoforms (Sepp et al. 2011). TCF-4A (short isoform) and TCF-4B (longest isoform) have been identified as the two main TCF-4 isoforms, yet their specific function is presently not understood. The isoforms of TCF-4 differ in their domain structure as the longest isoform contains an additional activation domain, the only nuclear localization signal and another repressor domain. In addition, analysis of transactivation efficiency has shown that TCF-4B has a higher capacity to induce gene expression than its shorter counterparts (Sepp et al. 2011). The present *Tcf4KO* mouse model displays residual expression of a short isoform of TCF-4 (TCF-4A), yet the fact that a loss of forebrain commissures was observed, strongly suggests that the longest isoform has singular functions in commissural formation. In this regard, future studies should compare the ability of TCF-4 isoforms for interaction with the identified transcription factor network and should map the respective interaction domains in the TCF-4 protein. An alternative, however, less likely explanation, given the additional functional domains of the long TCF-4B isoform would be that commissural development is highly dependent on TCF-4 dosage irrespective of the expressed isoforms.

Callosal abnormalities have been described in patients with PTHS who carry loss-of-function mutations in the bHLH domain (Amiel et al. 2007; Jung et al. 2018; Zweier et al. 2007). Disruption of the bHLH domain results in impaired DNA-binding affecting the transcriptional function of all TCF-4 isoforms (Sepp et al. 2012). Mutations in the first seven exons of *TCF4,* which do not affect the bHLH domain and may allow for the expression of shorter isoforms with an intact bHLH domain have been found in patients with moderate ID (Bedeschi et al. 2017). The present data raise the interesting possibility that such mutations may be sufficient to disrupt the function of TCF-4 in the development of interhemispheric connectivity and it would be interesting to investigate if these patients also display abnormalities in intercortical connectivity. In line with a previous study by Li and colleagues (Li et al. 2019), who analysed mice homozygously carrying a loss-of-function mutation affecting the bHLH domain of TCF-4, we found that loss of TCF-4 promotes the generation of SATB2+ neurons at the expense of deep layer neurons. While we have not analysed the molecular basis of these alterations, these data indicate that the loss of the commissural system in *Tcf4*KO mice is not the result of a failure to generate interhemispheric projection neurons and that TCF-4 is dispensable for the specification of SATB2+ neurons.

Dysplasia of the commissure system may not only be caused by impaired development of the respective projection neurons, but also by the failure to properly form a midline (Richards et al. 2004). In our analyses, the cellular composition of the midline appeared unaffected. At this point, we cannot fully exclude the possibility that subtle defects in midline cell composition and the erroneous display of axonal guidance cues contributed to the CC defects. The failed formation of the AC and the hippocampal commissure, though, hints at a general defect of neurons for commissure formation as these structures do not depend on midline fusion (Raybaud 2019).

While this study was in review, Mesman and colleagues reported the agenesis of the forebrain commissure system in a different *Tcf4* knockout mouse model (Mesman et al. 2020), which underlines the importance of TCF-4 in establishing interhemispheric connectivity. In contrast to our study, Mesman and colleagues found subtle defects in midline formation (Mesman et al. 2020). These phenotypic differences may be explained by the different genetic setup of the respective *Tcf4*KO models. While the present *Tcf4*KO model allowed for the residual expression of shorter TCF-4 isoforms with a functional bHLH domain, the *Tcf4* KO strain analysed by Mesman and colleagues harboured a mutation that removed the DNA binding bHLH domain from all isoforms (Mesman et al. 2020). Hence, future studies should address to what extent different TC-F4 isoforms contribute to the development of the midline.

Bulk RNA-Sequencing analyses of the developing murine neocortex of *Tcf4*KO mice showed that TCF-4 regulates a diverse set of genes with functions in cell proliferation, neuronal differentiation, and neurotransmitter release (Li et al. 2019; Mesman et al. 2020), which reflects TCF4’s pleiotropic functions in cortical development (Li et al. 2019; Mesman et al. 2020; Page et al. 2017). However, given the broad expression of TCF4 (Jung et al. 2018) and the considerable cellular diversity in the developing neocortex, bulk RNA-Sequencing data is not ideally suited to molecularly explain the complex phenotype of *Tcf4*KO mice and to identify cell-type and or stage-specific TCF-4 dependent mechanisms. In the present study, we used comparative single cell RNA-Sequencing analysis to zoom in onto the TCF-4-dependent transcriptome in *Satb2*-expressing neurons. We thereby uncovered that TCF-4 regulates genes with functions in axon guidance and neuronal and axonal development in this cell population, which provides a molecular explanation for the commissural phenotype in *Tcf4*KO mice. As the present data set contains single-cell RNA sequencing data from the developing neocortex, it provides an important resource to identify the cell type specific TCF-4 dependent transcriptome for other defined neocortical cell populations. Such analyses are expected to provide a better understanding of how TCF-4 functions as a pleiotropic regulator of cortex development.

Our approach of gene regulatory network analysis enabled us to identify how TCF-4 may affect these downstream targets. Classically, it had been assumed that TCF-4 partners with tissue specific bHLH TFs to influence transcription. Our results together with a recent *in vitro* study in mouse neuronal stem cells (Moen et al. 2017) indicate that TCF-4 also interacts with a multiplicity of TFs and chromatin remodelers outside the bHLH family to modulate their transcriptional activity. Here, we provided the first *in vitro* and *in vivo* biochemical evidence for these interactions and expanded the interactome of TCF-4 in a cell type specific manner for postmitotic intercortical projection neurons. As the scRNA data set is not restricted to intercortical projection neurons, it allows to predict cell type specific TCF-4-interactors also in other neocortical cell types, which will help to promote the understanding of how TCF-4 regulates the development of distinct cell populations.

Current evidence suggests that CC dysgenesis significantly contributes to cognitive impairment and associative dysfunction in intellectual disability, autism, and schizophrenia (Arnone et al. 2008; Badaruddin et al. 2007; Bedeschi et al. 2006; Hallak et al. 2007; Jeret et al. 1985; Paul et al. 2007; Rao et al. 2011; Siffredi et al. 2013). Intriguingly, many of the identified interactors are associated with these disorders and several of them are themselves linked to structural abnormalities of the CC (Cargnin et al. 2018; Filatova et al. 2019; Pinero et al. 2020; Pringsheim et al. 2019; Snijders Blok et al. 2019; Tzeng et al. 2014).

In-depth study of the interaction of TCF-4 with SOX11 - provided *in vivo* biochemical and functional evidence for cooperativity of neurodevelopmental disorder-linked genes in the generation of the commissural system and for the regulation of factors suggested in the pathogenesis of neuropsychiatric disease. We propose that the present data provides a new entry point towards understanding central dysregulated networks in the pathogenesis of autism and schizophrenia. Finally, we demonstrate that scRNA-Seq data can be harnessed to predict interaction partners of proteins. This powerful approach will be valuable to infer cell type specific transcription factor networks from complex tissues thereby enabling the discovery of regulatory networks in development, physiology and disease.

## Material and Methods

### Experimental models

All experiments were carried out in accordance with the European Communities Council Directive (86/609/EEC) and were approved by the government of Middle-Franconia. Tcf4ex4WT/lacZ mice were obtained from the Wellcome Trust Sanger Institute and previously described in Jung et al. (2018) (Alleles produced for the EUCOMM and EUCOMMTools projects by the Wellcome Trust Sanger Institute; MGI ID: 4432303). The Sox11^LacZ/WT^ mice were previously described (Sock et al. 2004).

Experiments were performed on male and female littermates between E16.5 and P56. For embryonic studies, mice were bred in the afternoon and vaginal post-coitum protein plug check (“Plug check”) was performed the next morning. This time point was defined as E0.5. Numbers of animals used in each experiment are indicated in the figure legends.

Genotyping of the mice was done using the following primers:

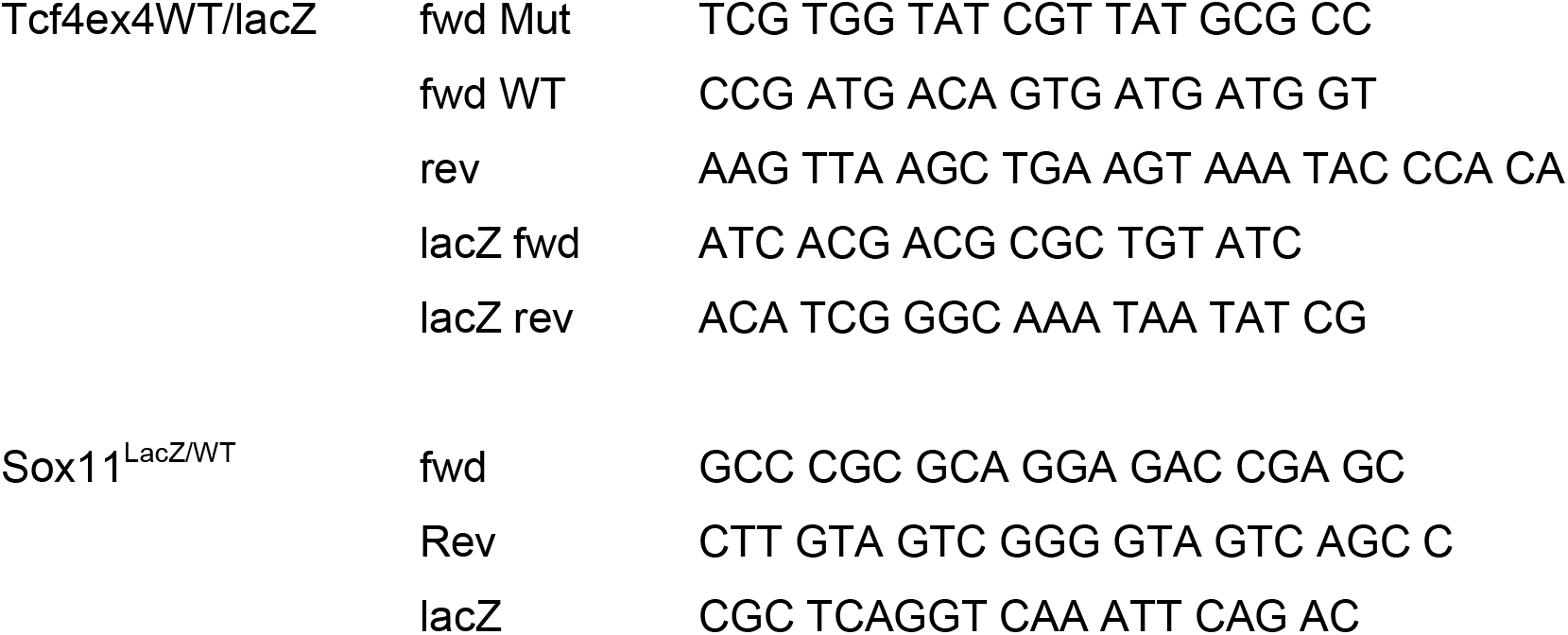

HEK 293T cells (ATCC, Wesel, Germany; CRL-3216) were in 10 cm dishes in DMEM supplemented with 10% of fetal bovine serum and 5 ml penicillin/Streptomycin at 37°C and 5% CO_2_.

### Experimental Design

For the single-cell RNA-Sequencing (5 WT and 4 KO samples) only samples with more than 500 cells after filtering were used to ensure a complete reproduction of cell diversity in the neocortex. Therefore, 2 samples for the WT and 2 samples for the KO were removed. We had to exclude one WT animal that displayed lower *Tcf4* expression than the KO and also excluded one cluster that displayed a high background transcript expression of blood related genes such as Hbb-a1, Hbb-a2.

### Tissue preparation and dissection

Timed pregnant mice were killed by cervical dislocation. For the E16.5, E18.5 and P0 time points, brains were dissected and fixed overnight in 4% PFA. Tails were used for genotyping. After fixation tissue was washed repeatedly with 1× PBS and transferred to 30% sucrose in 0.1 M phosphate buffer overnight for dehydration. Embryonic tissues were embedded in freezing media (Leica Biosystems, Richmond) and stored at − 80°C. Adult mice were killed using CO_2_ and transcardially perfused with PBS for 2 min (20 ml/min) followed by fixation with 4% paraformaldehyde (PFA) in PBS, pH 7.4, for 5 min. The brains were post-fixed overnight in 4% PFA at 4°C followed by dehydration at 4°C in 30% sucrose in 0.1M phosphate buffer.

### Histology

Embryonic tissue was cut in 10 μm thin sections with a cryotom (Leica Microsystems, Wetzlar). Sections were transferred on laminated object slides, dried for 2 h at room temperature and stored at − 80°C until further use. Slides were washed once for 5 min with 1x PBS. For antigen retrieval, sections were treated with 10 mM citrate buffer (pH 6) for 11 min at 720 watt in the microwave. Afterwards, half of the citrate buffer was replaced by water and the sections were incubated for another 30 min. Further steps were performed for both antigen retrieval and normal staining protocol. Slides were washed once in 1x PBS and subsequently incubated in 4% PFA for 10 min followed by two more washing steps in 1x PBS. Tissue was permeabilized for 10 min in 0.3% Triton-X/PBS and blocked with blocking solution (10% Donkey serum, 3% BSA and 0.1% Tween20 in PBS) for at least 1 h in a wet chamber at room temperature. Sections were incubated with primary antibodies [rb BRN1 (kind gift of Elisabeth Sock) 1:500); rb BRG1 (Santa Cruz, sc10768) 1:100; rb CALRETININ (Swant 7699/4) 1:500; rt CTIP2 (Abcam, 18465) 1:500; ab rb FOXG1 (Abcam, ab18259) 1:500; rb GAP43 (Abcam, ab5220) 1:500; ch GFAP (Abcam, ab4674) 1:500; ms SATB2 (Santa Cruz, sc-81376), 1:500; rat anti-SOX11 (kind gift from Johannes Glöckner) 1:500; rb TBR1 (Abcam, ab31940) 1:500; ms TCF-4 (Santa Cruz, sc393407) 1:100] diluted in blocking solution at 4°C overnight. Slides were washed three times for 5 min with 1×0.1% Tween/PBS, incubated with secondary antibodies diluted in blocking solution for 2 h at room temperature, and washed three times with 1x PBS. Nuclei were stained with DAPI (500 pg/ml in 1x PBS) for 10 min. After additional washing with 1x PBS for 5 min, slides were mounted with 60 μl Mowiol (Sigma Aldrich Chemie GmbH Munich, Germany) and stored at 4 C.

### Cell counting

Cell counting was done blind to avoid bias. Numbers were randomly assigned to slides before imaging. Genotypes were only revealed for statistical analysis. All images of the cortices were taken with the pial surface at the upper edge of the picture and the ventricular surface at the lower edge. Cells in an image were counted using ImageJ software and reported as the total numbers of cells per surface area of the VZ.

### Lipophilic Tracer Analysis

For the lipophilic tracer experiment P0 brains were dissected, washed once in 1xPBS and dried on a soft tissue. 1 μl of DiI dilution [DiIC18(3), Invitrogen, Eugene, Orgeon] was pipetted on one hemisphere and the brain subsequently fixed in 4% PFA. After six weeks the tissue was transferred to 30% sucrose in 0.1 M phosphate buffer overnight for dehydration, then frozen in tissue freezing media (Leica Biosystems, Richmond) and stored at − 80 C. Brains were cut in 10 μm thin sections with a cryotom (Leica Microsystems, Wetzlar). Sections were transferred on laminated object slides and dried for 2 h at room temperature. Slides were washed three times with 1x PBS and nuclei stained with DAPI (1:10.000 in 1xPBS) for 10 min. After additional washing with 1x PBS for 5 min, slides were mounted with 60μl Mowiol (Sigma Aldrich Chemie GmbH Munich, Germany) and stored at 4°C.

### Luciferase Assay

The ECR from the *Plxna2 gene* had the following positions in Mm10: chr1:194607209-194608066. The ECR was obtained by PCR from WT mouse DNA and inserted into the pTATA luciferase reporter plasmid in front of a β-globin minimal promoter. The hDCX-Promotor plasmid has been described before (Karl et al. 2005). HEK cells were seeded in a density of 80.000 cells per well in a 24 well plate and transfected the next day. 400 ng of CAG-GFP-based expression vectors (CAG-GFP; CAG-*Sox11*-IRES-GFP (Balta et al. 2018); CAG-TCF-4-IRES-GFP (Jung et al. 2018)), 200 ng of luciferase reporter (hDCX-pGL3 (Karl et al. 2005) and pTataLuc-Plxna2) and 10 ng of Renilla expression plasmid per well were transfected using JETPEI (Polyplus transfection, 101-10 N) according to the manufacturers’ instruction. Three wells were transfected per condition as technical replicates. After 48 h Luciferase assay was performed according to manufacturers’ instruction using the Dual-Luciferase Reporter Assay System Kit (Promega).

### Luxol fast blue staining

To stain for myelin with luxol fast blue (Polyscience, Hirschberg an der Bergstraße) free floating sections were washed two times with 1x PBS, mounted on coated adhesive glass slides and dried for at least 2 hours at RT. The glass slides were incubated in luxol fast blue solution at 57°C overnight and then washed one time in 95% ethanol and one time in distilled water. The staining was differentiated in lithium carbonate solution for 3 min followed by incubation in 70% ethanol till white and grey mater was distinguishable. If this takes longer than 5 min, the glass slides are washed in distilled water again and the differentiation steps are repeated until white and grey mater are distinguishable from each other. The nuclei were stained with Mayer’s hemalun solution for maximal 30 sec and excess solution was removed by rinsing with tap water. Slides were mounted with 60 μl Mowiol and stored at 4°C.

### Imaging

For overview images and cell counting, fluorescence signal was detected with an AF6000 Modular Systems Leica fluorescent microscope and documented with a SPOT-CCD camera and the Leica software LAS AF (Version 2.6.0.7266; Leica Microsystems, Wetzlar Germany). For the analysis of the Luxol fast blue staining, images were obtained with a Zeiss MN Imager and x 2.5 objective lens. For co-expression analysis, fluorescence signal was detected using a Zeiss LSM 780 confocal microscope with four lasers (405, 488, 550, and 633 nm) and × 40 objective lens. Images were processed using ImageJ.

### Co-Immunoprecipitation

For *in vitro* Co-Immunoprecipitation HEK 293T cells (ATCC, Wesel, Germany; CRL-3216) were seeded in a density of two million cells in 10 cm dishes in DMEM supplemented with 10% of fetal bovine serum and 5 ml penicillin/Streptomycin. At a confluency of 70-90% cells were transfected using JETPEI (Polyplus transfection, 101-10 N) with equal amounts of the expression vectors (7.5 μg/10 cm dish) of CAG-TCF-4-IRES-GFP and the predicted interaction partners [pCMV5 rBrn1(Schreiber et al. 1997); pBJ5-hBRG1 (pBJ5 hBRG1 was a gift from Jerry Crabtree (Addgene plasmid # 17873; http://n2t.net/addgene:17873; RRID:Addgene_17873))(Khavari et al. 1993); pXJ42-p200 CUX1 (pXJ42-p200 CUX1 was a gift from Alain Nepveu (Addgene plasmid # 100813; http://n2t.net/addgene:100813; RRID:Addgene_100813))(Wilson et al. 2009); CAG-*Foxg1*-IRES-RFP; CAG-*Sox11*-IRES-GFP(Balta et al. 2018)) according to the manufacturer instruction. After 48 h, cells were harvested in 1 ml Buffer A [10 mM Hepes, pH 7.9, 10 mM KCl, 0.1 mM EDTA, pH 8.0, 0.1 mM EGTA, pH 8.0, protease inhibitor EDTA free cocktail (Roche PVT GmbH Waiblingen, Germany) and Phosphatase Inhibitors Cocktail (Sigma Aldrich Chemie GmbH Munich, Germany)] (330μl per 10 cm dish). Three 10 cm dishes were combined for every experiment. After addition of 100 μl of 10% NP-40 and 84 μl of 5 M NaCl the solution was vortexed for 10 sec followed by 15 min of incubation on a rotating wheel at 4°C. The homogenates were centrifuged at 14000xg for 3 min. The supernatant was used directly for the co-immunoprecipitation by mixing 300 μl with 1.2 ml of TEN-Buffer [10 mM Tris, pH 7.4, 0.05 mM EDTA, 50 mM NaCl, 0.25% 10%NP40, protease inhibitor EDTA free cocktail (Roche PVT GmbH Waiblingen, Germany) and Phosphatase Inhibitors Cocktail (Sigma Aldrich Chemie GmbH Munich, Germany)] and 2 μl of ms TCF-4-antibody (Santa Cruz, sc393407), 2 μl of ms BrdU-antibody (BD Bioscience, B44) (Control for unspecific binding to mouse antibodies), or nothing to control for unspecific binding to Protein A Agarose Beads, Fast Flow (GE Healthcare Bio-Sciences AB, Uppsala, Sweden). An appropriate amount of the supernatant was kept as Input. Prepared probes were incubated on a rotating wheel at 4°C overnight. 30 μl of Protein A Agarose Beads, Fast Flow (Millipore-Merck, Darmstadt) in TEN-Buffer (1:1) were added and the samples were rotated for another 3 h at 4°C. Samples were centrifuged for 5 min at 1200xg and the supernatant was discarded. Beads were then washed three times with 500 μl of TEN-Buffer and frozen at −80°C. For Western Blot analysis 30 μl of 3xLaemmli buffer was added to the beads and incubated at 95°C for 5 min. 30 μl of the samples were loaded on 10% 1,5 mm SDS gels.

For *in vivo* Co-Immunoprecipitation, neocortices of E18.5 WT embryos were dissected and either used directly or stored at −80°C until further use. Two cortices were merged and homogenized in 1 ml of Buffer A. Samples were treated as described above. Antibodies used were rb SOX11-antibody (Abcam, ab134107) and rb EPHA3-antibody (Abcam, ab110465). For Western Blot analysis 50 μl of 3xLaemmli buffer was added to the beads and incubated by 95°C for 5 min. 50 μl of the samples were loaded on 10% 1,5 mm SDS gels.

### Western Blot

Protein extracted from E18.5 WT or KO cortices were obtained by homogenizing the tissue in RIPA buffer [50 mM Tris-HCl, pH 8.0, 150 mM NaCl, 1% Nonidet P-40, 0.5% Na-deoxycholate, 0.1% SDS, 2 mM EDTA, protease inhibitor EDTA free cocktail (Roche PVT GmbH Waiblingen, Germany) and Phosphatase Inhibitors Cocktail (Sigma Aldrich Chemie GmbH Munich, Germany)] followed by incubation for 30 min on ice. The post-nuclear supernatant of the lysate was obtained by centrifugation at 2000xg for 10 min at 4°C. Protein content was measured using the Pierce BCA protein assay (Thermo Scientific, Warrington, UK). For Western Blot analysis 30 μg of protein were loaded on a 10% 1mm SDS-PAGE gel. Gels underwent wet transfer onto a nitrocellulose membrane. Membranes were blocked in PBS with 0.1% Tween 20 (PBS-T). Incubation with primary antibodies [rb TCF-4 (Abcam; ab130014) 1:500] diluted in 5% BSA in PBS-T was performed overnight at 4°C and was followed by three times washing with PBS-T. Secondary antibodies were diluted in PBST and incubated with the membranes for at least 1 h at room temperature followed by washing with PBS-T. Membranes were treated with Clarity Western Enhanced Chemiluminescence Substrate (Bio-Rad) and visualized with Fusion-SL (PeqLab). Images were processed via Fusion (PeqLab).

### Single-cell RNA sequencing and analysis

#### Single-Cell Isolation of E18.5 cortex tissue

Neocortices of E18.5 embryos were dissected under a binocular. Each cortex was incubated in 150 μl of Ovomucoid-Mix [1.15 mg/ml Trypsin-Inhibitor (Sigma Aldrich Chemie GmbH Munich, Germany), 0.53 mg/ml BSA, 400 ng/ml DNase I Type IV (Roche PVT GmbH Waiblingen, Germany) in L15 medium (Gibco)] and carefully cut into small pieces. After addition of 150 μl of Papain-Mix (30 U/ml Papain (Sigma Aldrich Chemie GmbH Munich, Germany), 0.24 mg/ml Cysteine (Sigma Aldrich Chemie GmbH Munich, Germany), 40 μg/ml DNase I Type IV (Roche PVT GmbH Waiblingen, Germany)) samples were incubated for 15-20 min at 37°C. To dissociate the cells. 300 μl of Ovomucoid were added followed by a 5 min incubation at room temperature. The tissue was then triturated with fire-polished glass pipettes and transferred to 10 ml of L15 medium. To obtain the cells, the solution was centrifuged for 5 min at 90xg and about 9.5 ml of the supernatant was discarded. Cells were resuspended in the remaining media and strained through a cell strainer (Mesh size: 40 μm) to remove clumps. Cell density was determined using a Neubauer chamber. Libraries were prepared using the Chromium Controller and the Chromium Single Cell 3’ Reagent Kit v2 (10X Genomics, Pleasanton, CA). Single cell suspensions were diluted in nuclease-free water according to manufacturer instructions to obtain a targeted cell count of 5000. cDNA synthesis, barcoding, and library preparation were then carried out according to the manufacturers’ instructions. The libraries were sequenced on an Illumina HiSeq 2500 (Illumina, San Diego) with a read length of 26 bp for read 1 (cell barcode and unique molecule identifier (UMI)), 8 bp i7 index read (sample barcode), and 98 bp for read 2 (actual RNA read). Reads were first sequenced in the rapid run mode, allowing for fine-tuning of sample ratios in the following high-output run. Combining the data from both flow cells yielded approximately 200 M reads per mouse.

### Data Processing for scRNA-seq Analysis Using Cell Ranger and Seurat

The reads were de-multiplexed using Cell Ranger (version 2.1.1, 10X Genomics) mkfastq and read quality was assessed by FastQC (version 0.11.8, Babraham bioinformatics). For mapping the reads to the mm10 genome (10X Reference 2.1.0, GRCm38, Ensembl 84) and to identify single cells the standard Cell Ranger workflow was used. Common quality control measures for scRNA-seq (gene count per cell, UMI count per cell, percent of mitochondrial transcripts) were calculated using the Seurat R package (version 2.3.4) (Butler et al. 2018; Satija et al. 2015). The analyses were performed for genotypes and for each mouse individually. Quality control thresholds were set to 1,000-5,000 genes per cells, 1800-10000 UMIs and <6% of mitochondrial transcripts. Only samples with more than 500 cells after filtering were used to ensure a complete reproduction of cell diversity in the neocortex. 3 samples for WT and 2 samples for KO were used for further analysis. We had to exclude one WT animal that displayed lower *Tcf4* expression than the KO and we also excluded cells that displayed a high background transcript expression of blood related genes such as Hbb-a1, Hbb-a2.

### scRNA-Seq clustering and differential gene expression analysis using Seurat

Clustering of the cells was performed using the Seurat packages for R following the vignettes of the authors (Butler et al. 2018; Stuart et al. 2019). Cluster identity was defined using known marker expression for the different cell types. To extract SATB2 expressing glutamatergic cells, the data was subset by accepting only cells belonging to the intermediate progenitor, newborn neuron, deep layer and upper layer clusters with counts for SATB2 above 0. Differentially expressed genes between the two genotypes were determined using the MAST algorithm as implemented implementation in Seurat (Finak et al. 2015). GO-terms were identified with the Panther online tool (GO-Slim biological process and GO biological process complete) (http://www.pantherdb.org) (Mi et al. 2019).

### Gene regulatory network analysis by SCENIC

Assessment of gene regulatory networks (GRNs) was performed using the R package SCENIC (version 2) (Aibar et al. 2017). Only genes expressed in at least three cells were considered for analysis. The analysis was performed according to the packages vignettes. After gene regulatory networks defined, the networks were binarized. To that end a threshold was set at the mean of the area under the curve. In cells below the threshold the GRN was considered not active (OFF) whereas in cells above it was considered active (ON). As cells clustered apart according to the genotype, a list of GRNs was identified which where only active in one genotype. It was hypothesised that TCF-4 interacts with the heads of the GRNs and modulates their activity. To obtain a manageable list of candidate genes which might interact with TCF-4 the list of GRNs heads were analysed using the Panther online tool (Mi et al. 2019). Only genes associated with the GO terms neurogenesis/neuron differentiation and with at least an expression in 1/4 of the cells in the SATB2 cluster were chosen for validation. Common targets of TCF-4 and SOX11 were found by intersecting the list of DEGs from the *Satb2* cluster with the predicted targets of the Sox11 regulon. Disease association was determined by querying the list of differentially active regulons and common targets of TCF-4 and Sox11 in the DisGeNET database (https://disgenet.org)(Pinero et al. 2020).

## Statistical analysis

To determine statistical significance Mann-Whitney-U test was performed using the ggplot2 implementation of R (*, P ≤0.05; **, P ≤0.01, ***, P ≤0.001) if not otherwise indicated. n is indicated in the figure legends. Data is depicted as mean ± SD. To determine whether differences in luciferase activities (Figure 4B and C) were statistically significant, a two-tailed student’s t-test was performed using the ggplot2 implementation of R (*, P ≤0.05; **, P ≤0.01, ***, P ≤0.001). Data is depicted as mean ± SD. Results from independent transfections were treated as biological replicates.

## Data and code availability

The accession number for the single-cell RNA Sequencing of E18.5 neocortices is GEO: GSE147247.

## Acknowledgments

We thank Silvia Cappello, Michael Wegner and all members of the Institutes of Human Genetics and Biochemistry for helpful discussions.

This work was supported by the Deutsche Forschungsgemeinschaft (DFG, German Research Foundation) [Grant numbers 270949263/ GRK2162 and LI 858/ 9-1], by the Interdisciplinary Centre for Clinical Research Erlangen [Grant number E16 to D.C.L. and A.R.], and the Bavarian Research Network “ForINTER” to D.C.L..

M.T.W. is member of the research training group 2162 “Neurodevelopment and Vulnerability of the Central Nervous System” of the Deutsche Forschungsgemeinschaft (DFG GRK2162/1).

## Author contributions

Conceptualization, M.-T.W., D.C.L., A.R; Investigation, M.-T.W., P.K., A.B.E., Formal analysis, M.-T.W., P.K., A.B.E., D.C.L., A.R; Resources and Funding acquisition, D.C.L., A.R; Reagents, E.S.; Writing-Original draft, M.-T.W., D.C.L., A.R.; Writing-Review and Editing, M.-T.W., D.C.L., A.R.; Supervision: D.C.L., A.R.

## Supplement

**Supplemental Dataset 1. Differential expressed genes in the Satb2 cluster and GO term analysis. Related to Figure 2.**

**Supplemental Dataset 2. Differential expressed genes in the limited Satb2 cluster and GO term analysis. Related to Figure 2.**

**Supplemental Dataset 3. Differential active regulons in the Satb2 cluster. Related to Figure 3.**

**Supplemental Dataset 4. Differential active regulons in the limited Satb2 cluster. Related to Figure 3.**

**Supplemental Dataset 5. Overlap of differential expressed genes and the predicted Sox11 regulon in the Satb2 cluster and GO term analysis. Related to Figure 4.**

